# A pathogen-associated odorant induces fear-like response regulated by an olfactory receptor STR-211 in *Caenorhabditis elegans*

**DOI:** 10.64898/2026.08.07.743461

**Authors:** Anubhuti Dixit, Anisha Bhola, Avachi Azad, Tithi Thakur, Hina Bansal

## Abstract

Exposure to chemical cues released by predator or pathogen can evoke anxiety or fear responses in prey/host animals such as fight, flight or freeze both at behavioral and molecular levels. Freezing is a fundamental anxiety response when fighting or fleeing aren’t feasible. Despite the potential relevance of freezing as a stress-coping mechanism, its behavioral and molecular underpinnings are not understood yet. At molecular level danger cues are perceived by chemosensory receptors expressed in sensory neurons which may further regulate the animal’s behavioral responses(Ye et al., 2024){Citation}. 2-nonanone (2-NA) is one of the principal volatile organic compounds secreted by many pathogenic bacteria infecting *Caenorhabditis elegans* as well as humans and may signal danger to worms. Here, we show that olfactory exposure to threat-associated cue 2-NA induces a reversible fear-like freezing response characterized by immobility and halted feeding in *C. elegans*. With the application of *in silico* and behavioral approaches we showed that 2-NA is one of the ligands for an olfactory G-protein Coupled Receptor (GPCR) STR-211 and RNAi knockdown of the receptor leads to a defect in 2-NA induced avoidance behavior in worms. We next discovered that STR-211 is required for immediate behavioral changes in *C. elegans* during freezing response against 2-NA. The study proposes an environment relevant animal model to mimic human anxiety and fear-like behavior, along with the identification of one of the olfactory GPCRs mediating this behavior. The model may help in understanding the neuromolecular basis of freezing response in human anxiety, contributing towards treatment of mental health disorders.

## 1. Introduction

Anxiety and fear are evolutionarily conserved normal responses to potential or real threats. The threat signal which are known to induce fear responses include the presence of a predator, dead conspecifics, pathogens or chemicals released by them (Lin et al., 2025; Liu et al., 2018; Mutic et al., 2016; Oliveira et al., 2014). Animals rely on chemosensation to detect and respond to environmental cues that signal threat. Volatile compounds released by predator or pathogen may induce a fight-or-flight response by acting as a warning signal for potential danger (Baumbach et al., 2024; Bhola et al., 2026). Freezing, a type of fear response existing in most animals, is caused by the overwhelming presence of a fear stimulus where flight or fight are not possible. Though previous studies have indicated that threat-associated volatile compounds induce fear responses in animals, the behavioral and molecular aspects of the fear-induced freezing response are underexplored.

*Caenorhabditis elegans,* a soil-dwelling nematode, lives in a complex natural environment and uses chemosensation to locate food and avoid pathogenic bacteria and harmful chemicals (Dixit & Bhattacharya, 2021). Sensory perception of odorants such as volatile organic compounds (VOCs) released by pathogenic bacteria plays an essential role in shaping host defence strategies by enabling early detection and avoidance of harmful microbes. 2-nonanone (2-NA), a ketone produced by several bacterial species, including pathogenic strains such as *Pseudomonas* and *Serratia sp.* encountered by *C. elegans* in its natural habitat. It is one of the volatile signatures of foodborne pathogens infecting humans (Chen et al., 2017; Popova et al., 2014). Previous studies have demonstrated that 2-NA elicits robust avoidance behavior in *C. elegans* (Kimura et al., 2010; Troemel et al., 1997), suggesting that it functions as a warning cue indicative of harmful environmental conditions. This avoidance response represents an adaptive behavioral strategy that enhances survival by minimizing exposure to pathogenic threats. Indeed, previous work has demonstrated that 2-NA possess nematicidal properties and direct exposure to high concentrations of 2-NA causes mortality of worms (Deng et al., 2022), while moderate concentration causes decrease in life span and health span of *C. elegans* (Sarkar et al., 2024). This indicates that 2-NA exposure may mimic an important environmental threat cue for *C. elegans*.

Chemosensation in *C. elegans* is mediated primarily by a large family of G protein-coupled receptors (GPCRs), many of which are expressed in amphid sensory neurons. These receptors enable the worm to detect a wide range of chemical stimuli, including odor, chemicals, stressors, and pheromones, with high specificity. *C. elegans* has 16 pairs of chemosensory neurons located in amphid sensilla of the worm head region that play a major role in recognizing volatile and soluble odorants. The olfactory neurons AWA and AWC detect attractive chemicals such as diacetyl, isoamyl alcohol, and butanone (Bargmann et al., 1993; Troemel et al., 1995), while the repellents 2-NA, 1-undecene, and 1-octanol are recognized through the sensory neurons AWB and/or ASH (Chao et al., 2004; Troemel et al., 1997). The recognition and discrimination of odorants and sensory transduction follow similar principles in worms and humans (Yu et al., 2022). Though *C. elegans* responds to around 30 volatile chemicals, only a few odorants have been mapped to their cognate receptors. Moreover, functional evidence for the role of these receptors in regulating aversive odorants-induced behavior is lacking.

The present study is designed to understand the role of an aversive odorant, 2-NA, in inducing a fear response in *C. elegans* without causing mortality. We demonstrated that 2-NA odor exposure evokes a reversible freezing response in *C. elegans* characterized by a complete pause of locomotion and feeding. We identified a candidate olfactory receptor STR-211 for 2-NA by applying a combination of computational, genetic and behavioral approaches. With the help of molecular docking and simulation techniques, we showed that 2-NA has strong binding affinity for the receptor STR-211. Next, by utilizing RNAi knockdown, we have validated the functional role of *str-211* against known behavioral phenotypes caused by 2-NA such as chemotaxis, food avoidance, and aversion response in *C. elegans*. In the end, we showed that STR-211 regulates 2-NA induced freezing and immediate aftereffects of fear response. We propose an aversive odor induced fear model in *C. elegans* and one of the olfactory receptors regulating this response.

## 2. Materials and methods

### 2.1. *C. elegans* culture conditions

*C. elegans* were cultured at 20°C on nematode growth medium (NGM) plates on standard *E. coli* OP50 diet containing 100µg/ml streptomycin (Dixit et al., 2020). The Bristol N2 strain was used as wild type (WT) and the MAH677 (sid-1(qt9) V; sqIs71) strain was used for all RNAi experiments for neuron specific gene silencing. Young gravid adult stage worms were used for all experiments unless mentioned otherwise. All experiments were performed at 20°C. 2-NA and 1-octanol were purchased from Sisco Research Laboratories Pvt. Ltd. (SRL), India. 1-undecene and diacetyl were purchased from Sigma-Aldrich.

### 2.2. RNAi gene knockdown

The RNAi by feeding method was performed using *str-211* RNAi clone from the Ahringer library (Kamath et al., 2003). NGM agar plates containing 100 µg/ml ampicillin and 1 mM isopropyl-β-D-thiogalactoside (IPTG) were used for experiments. Bacterial cultures of control vector pL4440 and RNAi clone of *str-211* were grown for 16 hours in liquid Luria Broth medium with 100 µg/ml ampicillin. Cultures were grown for 1 hour after addition of IPTG (1 mM) and spread onto RNAi feeding plates. Plates were stored at room temperature for 24 hours before use. MAH677 worms were allowed to lay eggs on plates that contain the control vector alone or *E. coli* that expresses double-stranded RNA of the *str-211* gene. Age-synchronized worms thus obtained were used for all RNAi experiments.

### 2.4. Behavioral assays

**Chemotaxis assay:** To assess the preference of control and *str-211* RNAi knockdown worms towards 2-NA, chemotaxis assay was performed as per the protocol described previously (Bargmann et al., 1993). In brief, age synchronized worms were washed with S Basal buffer (100 mM NaCl, 50 mM K2HPO4 [pH 6]) thrice. 9 cm plates containing agar were divided into four equal quadrants. 2 μl of 2-NA (100% or 10%) or water was placed on each corner of the plate. Worms were placed at the center of the plate and will be allowed to move for one hour. 1 μl of 1M sodium azide was added after 1 hour, and the number of worms on each side were counted after 1 hour. Chemotaxis index was calculated according to the formula: (worms on odorant side)‒(worms on control side)/(total number of worms). Number of animals used for the experiment was between 50-60 per group and minimum three such independent experiments were performed.

#### Lawn avoidance assay

The lawn or food avoidance assay was performed as described previously with slight modifications (Harris et al., 2019). 50 μl of freshly prepared E. coli OP50 bacterial culture was seeded on 35 mm plate to create a 1 cm food lawn, and plates were kept at room temperature for 24 hours before use. Young gravid adult worms grown on control (vector) or *str-211* RNAi cultures were transferred to these plates and acclimatized for 60 mins. Next, a drop of 1 μl 100% 2-NA was placed on the right-hand side of the lawn. The number of worms inside and outside the lawn was counted every 5 minutes for a total of 15 minutes and the percentage of worms outside the lawn after 10 mins of the start of the experiment was plotted on the graph. Number of animals used for the experiment was 25 per group and three such biological replicates were performed.

#### Freezing response assessment

For assessment of threat induced freezing response, we have used 2-NA odor as the threat signal. We have exposed the worms to the odor of 2-NA by putting the chemical only on the lid, and the worms were placed on the plate. Total 3 μl of 100% 2-NA was placed on the lid by putting 1 drop of 1μl on three different points of an imaginary triangle (Figure 1a). We have made this arrangement so that the worms are exposed only to the odor of 2-NA which gets diffused all over the plate with no area available to worms for escape. Plates were tightly closed by parafilm and were observed under the stereomicroscope for around 3 minutes or till 80% of worms were immobilized. The number of worms immobilized after odor exposure were noted every 30 secs and the time taken for 100% of worms to immobilize was recorded for each group. Immediately after the worms became immobilized, the lid containing 2-NA was replaced with a blank lid to observe whether immobile worms were regaining their locomotion or not. Plates were observed at the intervals of 1 hour for a total of 2 hours to analyze the restoration of normal behavior of worms after 2-NA induced fear. A total of 50 animals were assessed for each group, and three such independent experiments were performed.

**Figure 1.**
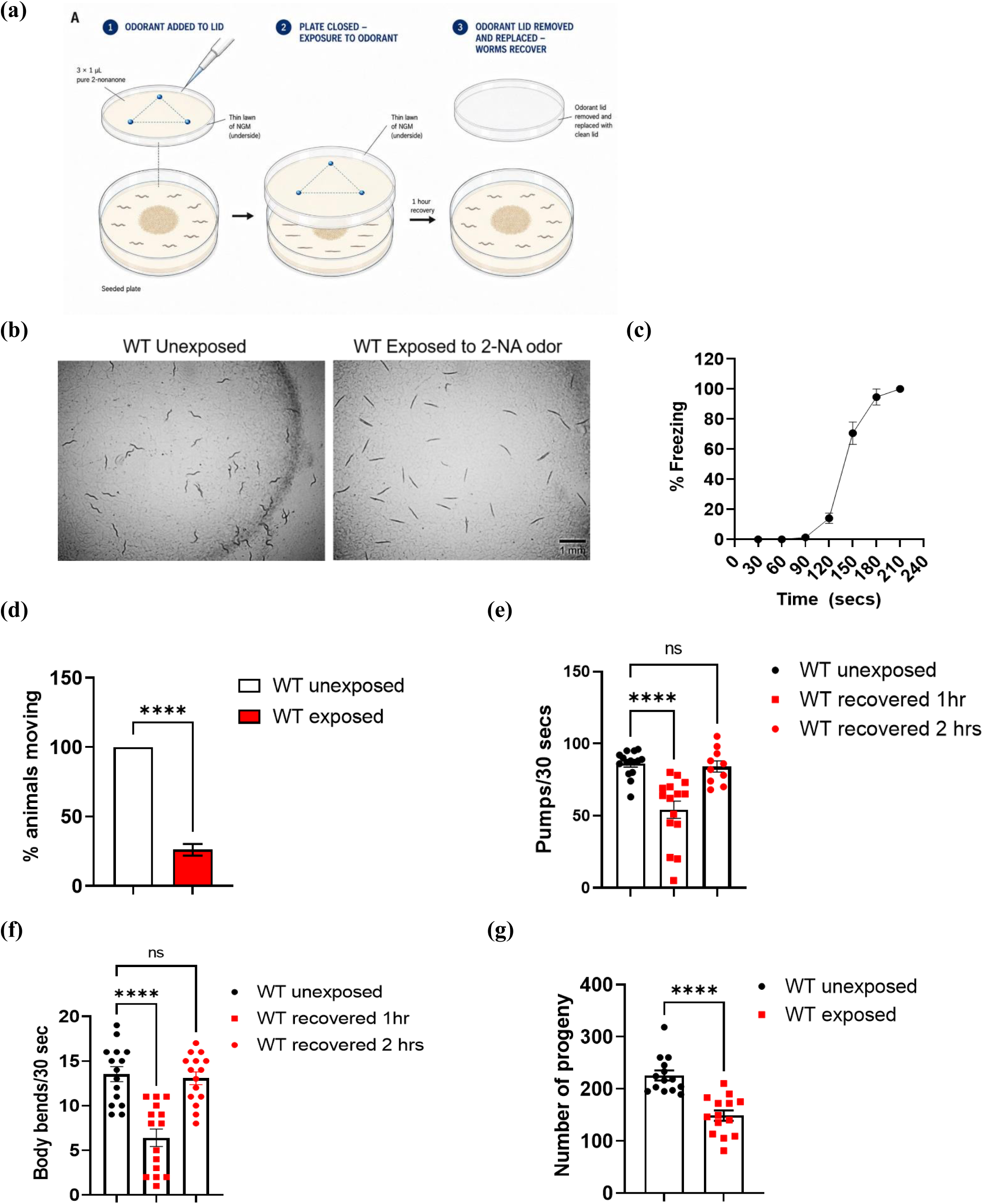
Exposure to 2-NA odor causes freezing of worms with cessation of locomotion and feeding in *C. elegans*. (a) Schematic diagram for freezing response assay by 2-NA odor exposure to worms, (b) Representative images of WT worms showing freezing response after exposure to 2-NA odor (scale bar 1 mM), (c) Percent animals showing movement when exposed to 2-NA odor for an average of 3 minutes, (d) feeding rates as measured by pharyngeal contractions and (e) locomotion of worms measured by number of body bends of WT worms before and after exposure of 2-NA odor. Measurements were taken after recovery period of 1 hour and 2 hours, (f) brood size of WT animals as measured by total number of progenies produced by each worm who have been exposed and not exposed to 2-NA induced freezing response. **** = p < 0.0001; ns= non-significant.

#### Feeding rate

The feeding rate was quantified by counting the pharyngeal pumping of synchronized well-fed worms. Pharyngeal pumping was assessed by manually counting the number of pharyngeal contractions of worms in 30-seconds under a stereomicroscope as reported earlier (Sarkar et al., 2024). A total number of 10-15 animals were assessed for each group.

#### Locomotion assay

Locomotion of worms was analyzed by counting the number of body bends in seeded NGM plates as reported elsewhere (Petratou et al., 2024). A complete body bend is when the whole head area, starting from directly behind the pharynx, moves to the opposite direction. 15-20 age synchronized worms were used for counting how many times the head of each animal changes direction per 30 secs for a period of 1 minute.

#### Aversion assay

The assay was done to analyse the acute reversal response of worms with sudden presentation of threat stimuli. We have done this assay to measure the response times of control and *str-211* knockdown worms to 2-NA as per the method reported previously with some modifications (Hilliard et al., 2002; Shukla et al., 2023). Young gravid adult worms were transferred to 35 mm

NGM plates without food. We have used a single worm per plate to quantify the reversal response. When a 2-NA drop is placed in proximity to the head of a worm, the worm senses it and reverses back, showing an aversion response (Figure 5a). The response was measured for 30 secs. Number of animals used for experiment was between 15-20.

### 2.3. Brood size measurement

Brood size was measured as reported elsewhere (Sarkar et al., 2024). Single late L4 stage worms of each control or *str-211* knockdown groups were exposed to 2-NA odor induced freezing for 4 minutes and individual worm was transferred to 35 mm plate seeded with respective culture, i.e. vector or *str-211* RNAi. The animals were transferred to a new plate every 24 hours till 4 days to avoid mixing day 1 progeny with day 2. Total brood size for each worm was determined by adding together the numbers of progeny produced by a single worm each day. A total number of 15 animals were assessed for each group.

### 2.5. *In silico* study

#### 2.5.1 Retrieval and Structural validation of selected proteins

We have selected three putative chemosensory GPCRs (STR-211, STR-92 and SRA-20) for our study, which were earlier shown to be defective in chemotaxis towards 2-NA (Taniguchi et al., 2014). The amino-acid sequence of Serpentine Receptor Class r-10 (STR-211), Serpentine receptor class alpha-20 (SRA-20), and Serpentine receptor class r-10 (STR-92) of C*aenorhabditis elegans* was retrieved from UniProt (UniProt ID: O16439), (UniProt ID: O17844), and (UniProt ID: O62279), respectively (https://www.uniprot.org/). Since no experimentally resolved three-dimensional structures were available for all three receptors, the AlphaFold-predicted structures were utilized for all computational analyses (Jumper et al., 2021; Varadi et al., 2022). The AlphaFold-predicted structures of all three were validated by the SAVES v6.0 server (https://saves.mbi.ucla.edu/). PROCHECK, ERRAT, and VERIFY3D performed structural quality assessment (Hashimoto et al., 1993; Kikuchi et al., 1993; Peplau, 1992). Ramachandran plot statistics, G-factor values, and overall quality scores were used to assess stereochemical reliability prior to molecular docking studies (Kikuchi et al., 1993).

#### 2.5.2 Binding Pocket Identification

The possible ligand-binding pockets of the three selected chemosensory receptors were predicted by using the PrankWeb (https://prankweb.cz/) and CASTp (https://cfold.bme.uic.edu/castpfold/) servers (Jendele et al., 2019; Ye et al., 2024). For STR-211, the top-ranked binding pocket predicted by PrankWeb had a pocket score of 31.64 and a probability of 0.899. The CASTp analysis also proved the existence of an outstanding cavity with a solvent accessible surface area of 318.973 Å2 and a volume of 157.473 Å3 Similarly, the predicted binding pocket of STR-92 had a PrankWeb score of 39.17, a probability of 0.936, and 22 pocket-lining residues. For SRA-20, the chosen binding pocket was detected by PrankWeb with a score of 3.59, a probability of 0.117, and 11 pocket-lining residues. The top-ranked or selected ligand-binding pockets found for each receptor were used to define the docking grids for all molecular docking analyses.

#### 2.5.3 Ligand Preparation

The odorant molecule was retrieved from the PubChem database: 2-NA (CID: 13187). The structure of the ligand was downloaded from the PubChem-NCBI database (https://pubchem.ncbi.nlm.nih.gov/) (Steinberg et al., 2023) in SDF format and converted to PDBQT format using OpenBabel (O’Boyle et al., 2011). The ligand energy was minimised and geometry optimised before docking.

#### 2.5.4 Molecular Docking

The binding affinity of the odorant molecule towards all three receptors was determined by molecular docking using AutoDock Vina (Trott & Olson, 2010). Docking calculations were carried out with a grid box of 40 × 40 × 40 Å centred on the predicted ligand-binding pocket of each receptor. The binding pocket prediction analysis was used to set the grid centers at x = 0.450, y = −2.312, z = −11.115 for STR-211, x = 5.4179, y = 4.6722, z = −3.2418 for STR-92, and x = −13.6055, y = −7.2022, z = 20.9794 for SRA-20. Ten docking poses were created for each receptor–ligand pair, and the highest-ranked conformation was chosen according to binding affinity and interaction profile.

#### 2.5.5 Molecular Dynamics Simulation

Molecular Dynamics Simulation of the docked STR-211+2-NA complex exhibiting the most favorable binding conformation was performed to understand the stability and conformational behaviour of the protein-ligand complexes using GROMACS (Kitami et al., 2016). This method enables the observation of atomic-level motions as a function of time and provides information about the stability, flexibility, and interactions of the docked complexes. The topology file of the target protein was generated using the CHARMM27 force field by means of pdb2gmx, and the topology and parameter files of the ligand were generated by the SwissParam server (https://www.swissparam.ch/) (Bugnon et al., 2023) to ensure compatibility with the CHARMM force field. The protein-ligand complexes were solvated in dodecahedron boxes by filling them with the TIP3P water model to simulate the explicit water environment.

Appropriate counterions were added to neutralize the simulation box. Following energy minimization, the system was equilibrated under NVT and NPT ensembles. A production molecular dynamics simulation was then performed for 300 ns. The trajectories were analyzed for the evaluation of important parameters like root mean square deviation (RMSD). These analyses provided an insight into the stability, flexibility and compactness of the protein-ligand complexes over the course of the simulation.

### 2.6. Statistical analysis

Data are presented as mean ± SEM. p < 0.05 was considered significant. Data was analyzed by one-way ANOVA when three or more groups are compared and by Student t-test when comparison was between two groups. The statistical software used for data analyses was Graph Pad Prism 9.0.

## 3. Results

### 3.1. Brief exposure to a pathogen-associated odor 2-NA induces fear-like freezing response in *C. elegans*

Previous studies have shown that when exposed to the volatile odorant 2-NA, worms show avoidance behavior. 2-NA is one of the VOCs secreted by many pathogenic bacteria that infect humans and worms. 2-NA displays nematicidal properties when worms come in direct contact with the chemical. *C. elegans* avoids 2-NA as it represents a threat to survival and acts as a danger signal to worms. We hypothesized that short duration indirect exposure to 2-NA may induce fear-like behavior in *C. elegans* without killing the worms. Therefore, we exposed the worms to the odor of 2-NA by placing the chemical on the underside of the lid of the plate (Figure 1a). We observed that brief (3-4 minutes) exposure to the odor of 100% 2-NA induces worms to undergo an immobile freezing state (Figure 1b and c). Immediately after the exposure to the odor of 2-NA, worms keep moving and searching continuously for the initial 90 seconds. In the next 30-seconds worms slow down, and between 120 to180 seconds from the time of 2-NA exposure, worms stop locomotion and feeding completely and undergo an immobile frozen state (Supplementary Video 1). This behavior could be a part of an innate fear response as found in most higher animals. We observed that increasing the duration of 2-NA odor exposure beyond 5 minutes results in the death of few worms. Therefore, we have strictly monitored the duration of exposure by keeping a stopwatch and removed the odor source immediately after all worms achieved a freezing state. Interestingly, the 2-NA induced freezing response was reversible and worms started regaining their movement and feeding gradually after odor source removal (Supplementary Video 1). Feeding rate and locomotion of worms returned to around 60% of normal level after 1 hour of recovery period. After 2 hours of recovery period both parameters completely restored to normal level as observed for unexposed worms (Figure 1d, e).

We have exposed worms to the odor of other pathogenic bacteria-associated volatile compounds such as 1-undecene and 1-octanol, which are known to induce repulsive behavior in *C. elegans* (Prakash et al., 2021; Troemel et al., 1997). We found all worms in moving condition after 4 minutes of odor exposure of equal volume (3 ul undiluted) of both chemicals (Figure S1a). However, when we have doubled the concentration and duration of odor exposure as 6 ul and 8 minutes, respectively, animals exposed to 1-octanol started undergoing a freezing state between 7-8 minutes of odor exposure (Figure S1b). The recovery from freezing state was slow in 1-octanol exposed animals, and they took around 30 mins to show locomotion again as compared to 4-5 minutes in the case of 2-NA. Therefore, we proceeded with 2-NA owing to its uniqueness for being recognized by worms as an immediate danger signal, which consequently induces shock and a faster fear-like response in animals.

Predatory cues induced fear response is known to decrease egg laying in *C. elegans* (Liu et al., 2018). We have previously shown that long-term exposure to a low dose of 2-NA decreases the brood size of worms (Sarkar et al., 2024). Therefore, we tested the number of progenies laid by worms who have experienced the 2-NA induced freezing response once before egg laying stage. We found that 3 minutes of exposure to 2-NA odor before the reproductive stage reduces the total brood size of worms upto 33% (Figure 1f). These results indicate that exposure to pathogen associated odorant 2-NA signals a potential danger to the worm and consequent fear-like freezing response. As soon as the danger signal is removed, worms return to their normal state. However, a single exposure of aversive odor stress creates a long-lasting impact on the animal’s reproductive capacity, reflected by reduction in total number of progenies produced by exposed worms.

### 3.2. Identification of candidate olfactory GPCR for 2-NA recognition

We next wanted to identify the molecular mediators of 2-NA induced avoidance and fear response in *C. elegans*. A previous large-scale RNAi screen to identify *C. elegans* olfactory receptors essential for the response to specific odorants has revealed three putative odorant receptors, STR-92, SRA-20, and STR-211 for the repellent 2-NA (Taniguchi et al., 2014). These receptors are expressed in various amphid neurons and belongs to the serpentine receptor family, a subgroup of GPCRs that are often associated with olfactory and chemosensory functions (Robertson & Thomas, 2006). Therefore, we screened all three GPCRs by performing molecular docking and simulation tools against 2-NA.

#### 3.2.1. Structural Validation of three putative odorant receptors

The structural evaluation of AlphaFold-predicted models of three selected receptors is shown in Figure 2. The analysis demonstrated high model quality for all receptors. Among them, STR-211 exhibited the highest structural reliability, with an ERRAT score of 99.70, followed by STR-92 and SRA-20, which achieved ERRAT quality factors of 93.88 and 93.66, respectively, indicating robust model accuracy. Ramachandran plot analysis further supported the structural integrity of the models, with 95.6%, 94.9%, and 96.2% of residues located within the most favoured regions for STR-211, STR-92, and SRA-20, respectively. No residues were observed in disallowed regions for STR-92 and SRA-20, while only 0.3% of residues in STR-211 fell within disallowed regions. The validation statistics are summarised in Supplementary Table S1.

**Figure 2.**
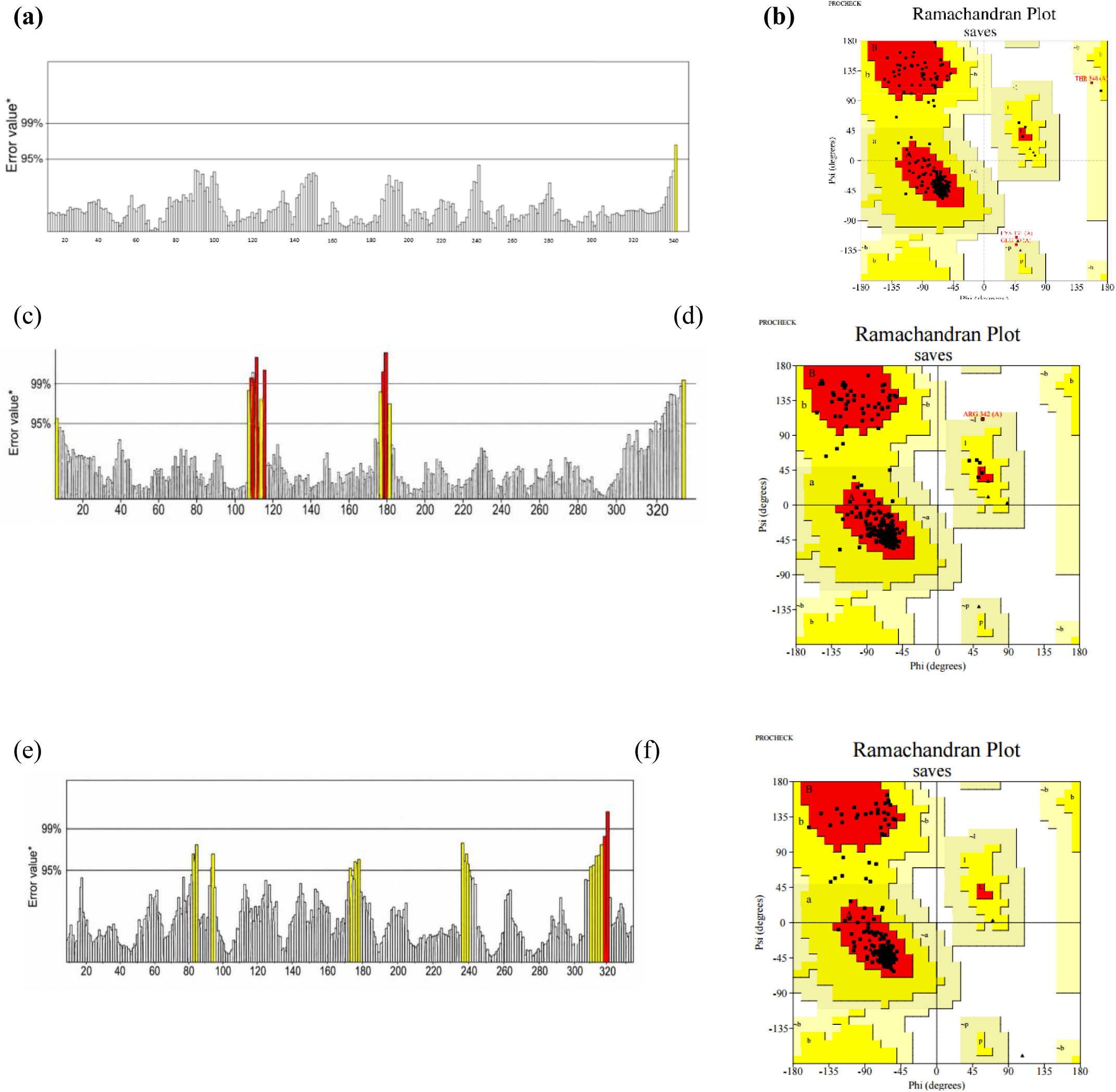
Structural validation of the AlphaFold-predicted chemosensory receptors. (a) ERRAT analysis of STR-211 showing an overall quality factor of 99.70. (b) Ramachandran plot of STR-211 showing 95.6% residues in the most favoured regions. (c) ERRAT analysis of STR-92 with an overall quality factor of 93.88. (d) Ramachandran plot of STR-92 showing 94.9% residues in the most favoured regions and no residues in disallowed regions. (E) ERRAT analysis of SRA-20 with an overall quality factor of 93.66. (f) Ramachandran plot of SRA-20 showing 96.2% residues in the most favoured regions and no residues in disallowed regions.

#### 3.2.2. Binding Pocket Identification

Potential ligand-binding cavities of the three receptors were predicted by PrankWeb (Jendele et al., 2019) and CASTp (Jin et al., 2026). Among the receptors, STR-211 exhibited a well-defined binding pocket (score: 31.64, probability: 0.899), which was further verified by CASTp analysis showing a comparatively large cavity. STR-92 demonstrated the highest prediction pocket confidence (score: 39.17, probability: 0.936), with the binding site characterised primarily by hydrophobic interactions. In contrast, SRA-20 showed a smaller and less confident binding pocket (score: 3.59, probability: 0.117), stabilised by hydrogen bonding and hydrophobic interactions. Variation in pocket size and architecture of the three receptors may be responsible for the difference in docking affinities observed in subsequent analysis. The predicted binding pocket and three-dimensional structural model of the protein receptors are shown in Figure 3, and their characteristics are summarised in Supplementary Table S2.

**Figure 3.**
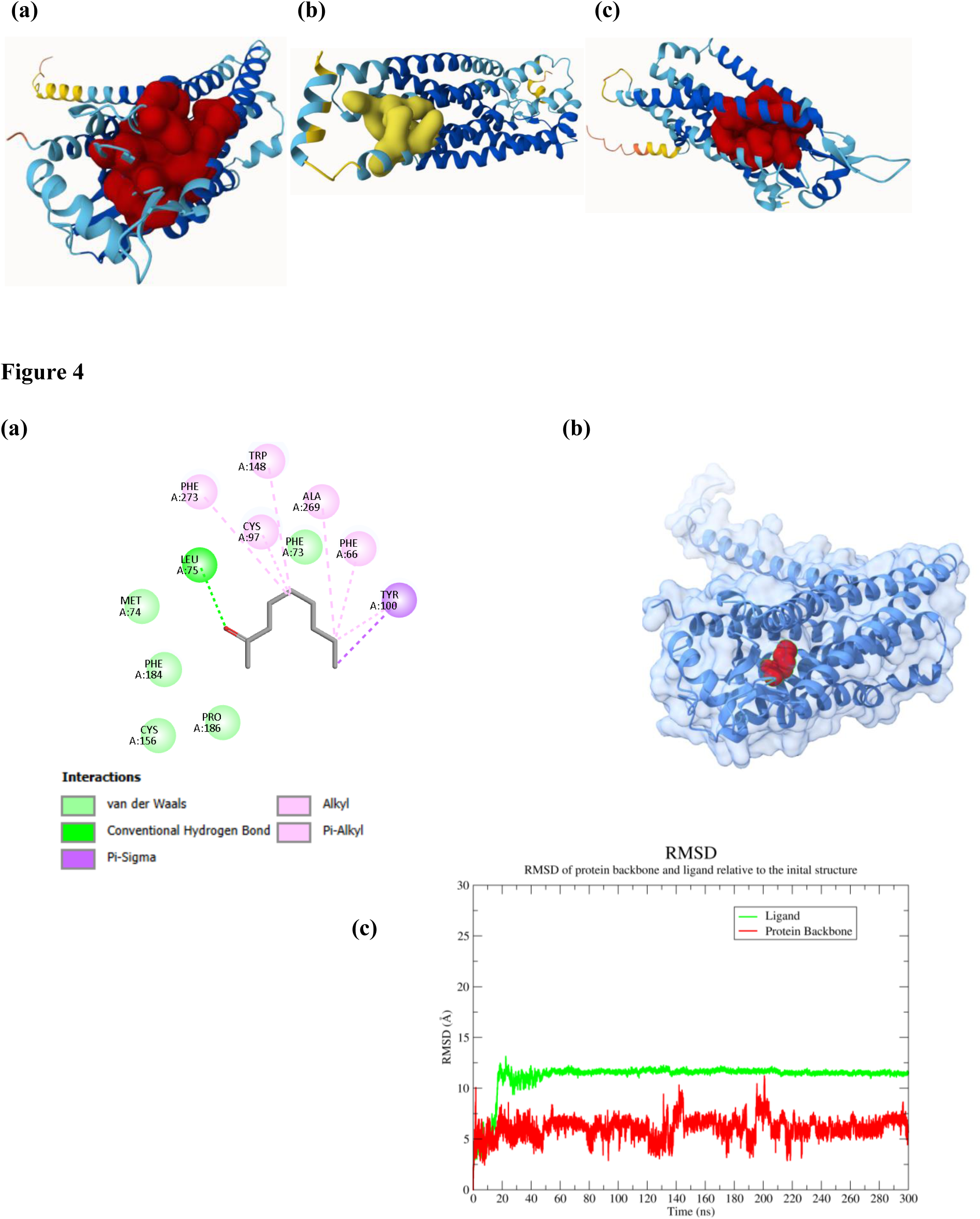
Predicted ligand-binding pockets of the AlphaFold-generated chemosensory receptors. (a) STR-211, (b) STR-92, and (c) SRA-20. The highest-ranked binding pockets predicted by PrankWeb and validated by CASTp are highlighted and were used to define the docking search space for subsequent molecular docking analyses.

#### 3.2.3. Molecular Docking Analysis and Molecular Dynamics Simulation

Molecular docking was performed to investigate the interaction of odorant molecule 2-NA with the three putative chemosensory receptors STR-211, STR-92 and SRA-20 of Caenorhabditis elegans. The docking results showed a significant binding score with all three receptors. Among the receptors examined, STR-211 showed the highest binding affinity (−5.2 kcal/mol), followed by STR-92 (−4.8 kcal/mol) and SRA-20 (−3.9 kcal/mol) (**Table 1)**.

**Table 1.** Comparative analysis of binding affinity of 2-NA with selected receptors.

| Receptor | UniProt ID | Binding Affinity (kcal/mol) |
| --- | --- | --- |
| STR-211 | O16439 | −5.2 |
| STR-92 | O62279 | −4.8 |
| SRA-20 | O17844 | −3.9 |

As the docking analysis indicated that STR-211 had the strongest affinity for 2-NA, this receptor–ligand complex was selected for a more detailed investigation of the molecular interactions and simulation analysis. We tested chemotaxis of worms against two different concentrations of 2-NA, i.e. 100% and 10%. We found that animals with str-211 knockdown were defective in chemotaxis behavior against all tested concentrations of 2-NA (Figure S2).

2D structural analysis of the docking complex revealed the formation of a conventional hydrogen bond with the residue Leu75, together with alkyl, π-alkyl, π-sigma, and van der Waals interactions with residues Phe66, Phe73, Tyr100, Trp148, Ala269, and Phe273. Notably, the binding cavity is enriched with aromatic and hydrophobic amino acids, suggesting that hydrophobic interactions are the primary contributors to ligand stabilization (Figure 4). The 2D interaction profiles of STR-92 and SRA-20 complexes are provided in Supplementary Figure S3.

**Figure 4.** (a) Two-dimensional interaction map illustrating the specific protein–ligand contacts observed in the docked complex, including hydrogen bonding, alkyl, π-sigma, π-alkyl, and van der Waals interactions. (b) Three-dimensional docking complex structure of the receptor STR-211 with the ligand molecule 2-NA. **(c)** Backbone RMSD of STR-211 and ligand RMSD of 2-NA during the 300 ns molecular dynamics simulation. The protein shows stabilization in beginning phase of equilibration and the ligand shows early conformational adaptation and then rather stable trajectory during simulation.

To evaluate the dynamic stability of the docked complex, a 300 ns molecular dynamics simulation was performed using GROMACS with the CHARMM27 force field and TIP3P water model. The RMSD profile of the protein backbone indicated an initial equilibration during the first 20-30 ns, followed by stabilization for the remainder of the simulation. The backbone RMSD values were mostly within the 5–7 Å range, indicating that the receptor retained its structural integrity, even though conformational adjustment was made due to the solvent equilibration.

The ligand RMSD analysis revealed a rapid increase in the early phase of the simulation that was related to the reorientation of 2-NA within the binding cavity. After this adaptation period, the ligand RMSD remained steady at around 11-12 Å and was relatively stable throughout the rest of the trajectory. The ligand RMSD reached a relatively stable plateau after an initial conformational adjustment, indicative of adaptation to a new energetically favourable binding orientation inside the receptor cavity. The transient fluctuations of 130-150 ns and 190-210 ns are most likely due to local conformational rearrangements, rather than a large-scale destabilisation. Overall, the RMSD analysis supports the structural stability of the STR-211+2-NA complex and validates the docking-predicted binding mode under dynamic conditions (Figure 4).

### 3.3. RNAi knockdown of *str-211* causes defects in avoidance behavior of worms against 2-NA

To understand whether the *str-211* receptor is required in *C. elegans* for repulsive response against 2-NA, we have conducted two behavioral tests; aversion assay and lawn avoidance. The aversion assay measures avoidance response in *C. elegans* against repellent. It is a rapid escape behavior where the nematode immediately stops forward movement, quickly backs up, and turns to navigate away from danger. It is a vital survival reflex triggered by sudden noxious stimuli. We tested the avoidance response of individual worm when exposed to 100% 2-NA. Avoidance behavior of *C. elegans* for 2-NA is regulated by transitions between the two behavioral states, pirouette and run. A pirouette is a period of short movements interrupted by frequent reversals and turns, and run is a period of long straight migration of >14 secs (Yamazoe-Umemoto et al., 2015). *C. elegans* efficiently chooses the appropriate migratory direction by application of these movement strategies (Figure 5a; Tanimoto et al., 2017). We found that when a drop of 2-NA is put on the path of control worms, they immediately turn back and undergo a brief period of pirouette state before initiating a straight run. However, *str-211* knockdown worms display a 21% increase in the duration of pirouette state than control worms (Figure 5b). The number of turns during the pirouette state were also more in *str-211* knockdown animals in comparison to control worms, indicating a defect in decision making to move away from harmful 2-NA stimuli (Figure 5c).

Next, we tried to understand whether STR-211 is also required for avoidance response against 2-NA when the odorant exposure is coupled with the presence of food. Previously, it has been shown that worms leave a food lawn when exposed to 2-NA simultaneously (Harris et al., 2019). Indeed, we observed that after 10 mins of exposure to 100% 2-NA near the bacterial lawn, around 65% of worms leave the lawn and reach to opposite direction of the plate. However, in the *str-211* RNAi group, worms are slow in leaving the lawn, and only 45% of worms are present outside the lawn after 10 mins of exposure to 2-NA (Figure 5d and 5e). The lawn occupancy of control and *str-211* RNAi worms is similar when not exposed to 2-NA (Figure S3). These experiments indicate a decreased ability of *str-211* knockdown animals to recognize 2-NA and to initiate a faster avoidance response in the absence or presence of food.

### 3.4. STR-211 is required for 2-NA induced fear-like response in *C. elegans*

We have shown that STR-211 receptor is involved in the recognition of 2-NA and regulates avoidance behavior against 2-NA, we planned to test the effect of *str-211* knockdown on 2-NA induced fear response in *C. elegans*. Results indicated that *str-211* RNAi animals exhibited a significant increase in the number of animals moving in comparison to the control group after 3 minutes of odor exposure of 100% NA (Figure 6a). The *str-211* animals also display faster recovery in pumping rate and locomotion of animals from freezing state after 1 hour of 2-NA odor exposure (Figure 6b and 6c). It shows that STR-211 receptor is required in *C. elegans* for induction of fear response by 2-NA. However, knockdown of *str-211* does not change 2-NA induced decrease in brood size of worms (Figure 6d), suggesting a role of the STR-211 receptor in regulating immediate fear responses against 2-NA not the long-term health effects caused by the odorant stress.

**Figure 5.**
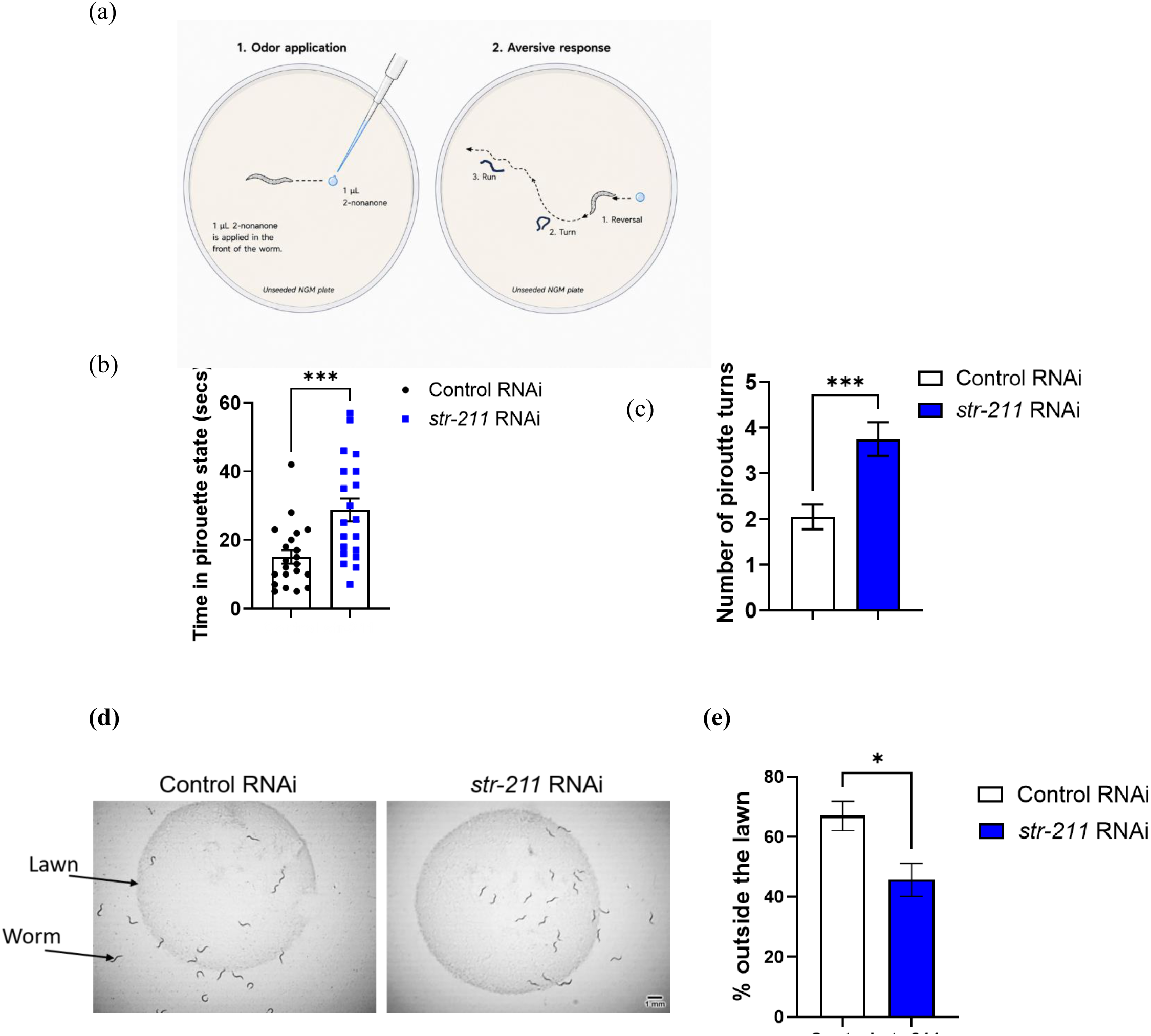
STR-211 regulates 2-NA avoidance behavior of *C. elegans*. (a) Schematic of avoidance assay showing reversal response of worms composed of pirouette turns and run state, (b) time spent by control and *str-211* RNAi animals in pirouette state before initiating run after encountering 2-NA on the path, (c) number of pirouette turns by control and *str-211* RNAi animals before initiating run after encountering 2-NA on the path, (d) Representative images and (f) graph for number of animals outside the lawn of control and *str-211* RNAi groups (scale bar 1 mM). *** = p < 0.001; * = p < 0.05.

**Figure 6.**
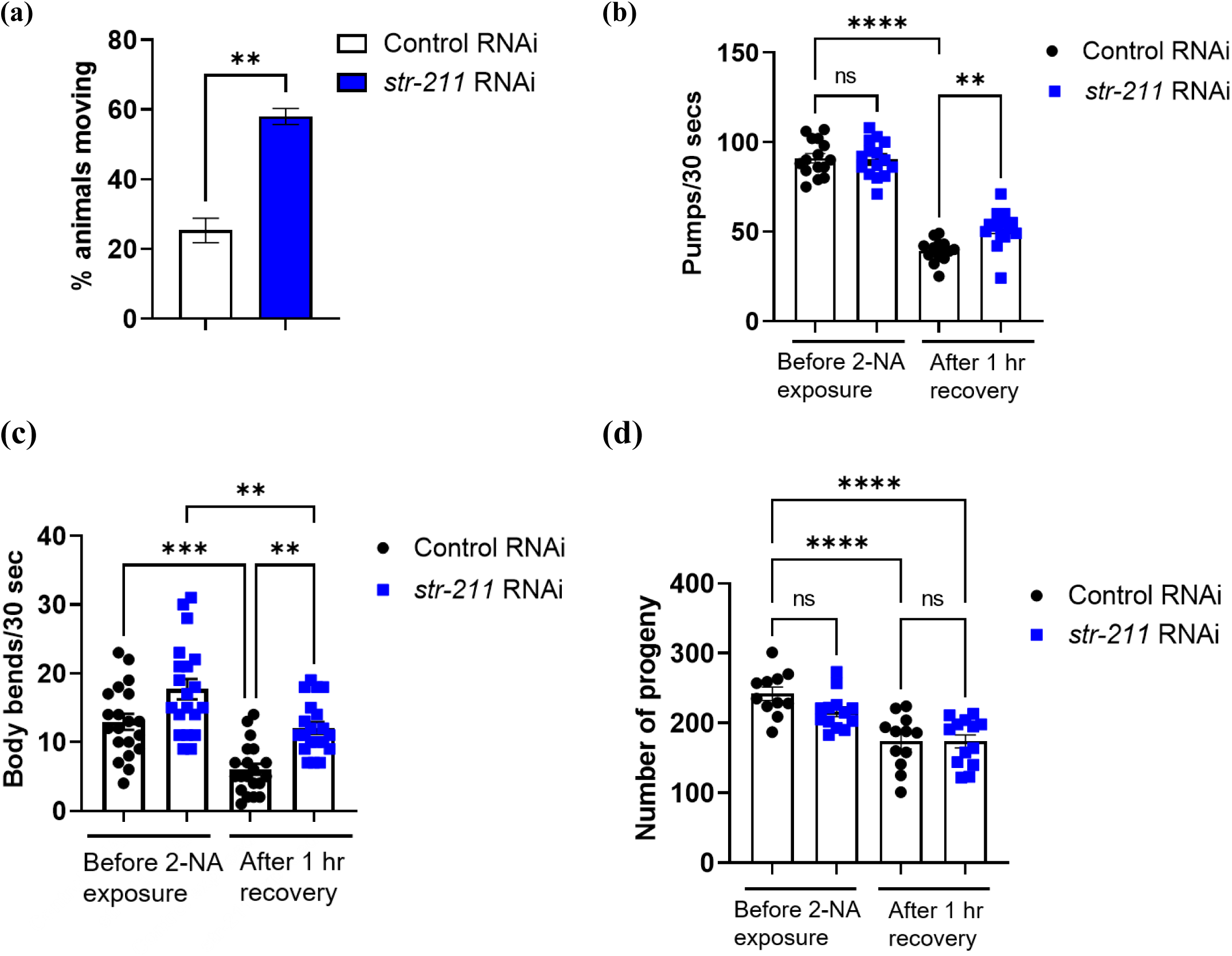
2-NA evoked fear-like behavior is mediated by olfactory receptor STR-211 in *C. elegans*. (a) Percent of control and *str-211* RNAi animals showing movement when exposed to 2-NA odor for an average of 3 minutes, (b) feeding rates as measured by pharyngeal contractions and (c) locomotion of worms measured by number of body bends of control and *str-211* RNAi animals before and after exposure of 2-NA odor. *str-211* knockdown animals display faster recovery in feeding and locomotion after 1 hour of 2-NA induced freezing response, (d) brood size of control and *str-211* RNAi animals as measured by total number of progenies produced by each worm who have been exposed and not exposed to 2-NA induced freezing response. **** = p < 0.0001; ** = p < 0.01.

Taken together, our findings demonstrate that a single brief exposure to pathogen-associated odor 2-NA in *C. elegans* induces a fear-like freezing response which alleviates after removal of the odor source. Knockdown of chemosensory receptor STR-211 makes worms slow in avoiding 2-NA and less susceptible to the freezing effect.

## 4. Discussion

Our study proposes a novel model of 2-NA odor-induced fear-like freezing behavior in *C. elegans*. Freezing is an innate fear response of the body when the nervous system feels overwhelmed by perceived threat. The freezing fear response is considered as an evolutionary conserved survival mechanism where the nervous system completely halts movement to assess a threat or avoid detection.

*C. elegans* perceive pathogens or associated compounds as potential threats and display an innate fight-or-flight response against them. As 2-NA is one of the volatile compounds present in various bacteria known to infect and kill *C. elegans* (Chen et al., 2017; Popova et al., 2014), worms sense danger when exposed to 2-NA odor and show a freezing reaction. Complete shutdown of feeding and locomotion in *C. elegans* during the freezing response reflects conservation of energy, which can be directed to increase the survival duration of animals. Our results emphasize the unique role of 2-NA odor in evoking a fast and robust freezing response which we could not find with other repellents. As reported for predator cue induced freezing behavior in rodents (Tyler et al., 2022), the unconditioned freezing response of worms against 2-NA is transient and reversible. However, fear induced stress has been implicated in producing long-lasting changes at behavioral and molecular levels in both human and animals (Pitman et al., 2012; Rodrigues et al., 2009). A recent study has shown that a single exposure to the predator odor 2,4,5-trimethylthiazoline to mice increases general anxiety even after 10 days of exposure (Baumbach et al., 2024). In our study, we observed that a single exposure to 2-NA odor causes a 1.5-fold decrease in the number of progenies produced over a period of 4 days by WT worms. This indicates a lasting effect of 2-NA odor-induced stress and can be compared with reduced fecundity in women suffering from posttraumatic stress disorder (PTSD) (Wamser-Nanney, 2020).

Our research shows that the volatile chemical 2-NA holds an ecological and functional significance for *C. elegans*. We decided to understand the downstream signalling mechanism for the effect of 2-NA odorant. Olfactory GPCRs are the first site of interaction between external odor cues and the internal cellular environment. We have shown by *in silico* and behavioral analysis that STR-211 is one of the olfactory receptors for 2-NA recognition in *C. elegans*. The AlphaFold-predicted structures of STR-211, STR-92 and SRA-20 displayed good stereochemical quality (Jumper et al., 2021; Varadi et al., 2022), which supported their use for structural and docking analyses. Comparison of binding pockets showed different cavity architectures (Jendele et al., 2019; Ye et al., 2024), which were also apparent in their docking behaviour. STR-211 showed the best binding affinity towards 2-NA, which we attribute to its well-formed binding pocket, hydrogen bond formation, and extensive hydrophobic interactions (Senior et al., 2020) in comparison to STR-92 and SRA-20, which formed relatively fewer stabilising contacts. To confirm the stability of the STR-211-2-nonanone complex, a 300 ns molecular dynamics simulation was carried out, which confirmed the reliability of the predicted binding mode under dynamic conditions. Our work indicates that binding of the repellent 2-NA with STR-211 is stronger (binding affinity -5.2 kcal/mol) than the binding of attractant diacetyl with ODR-10 (binding affinity around -4 kcal/mol), a well-established odor-receptor pair. Similar to recognition of diacetyl by ODR-10, we showed the dominance of hydrophobic interactions for molecular recognition of 2-NA by STR-211 in protein ligand interaction analysis (Di Rienzo et al., 2026). Moreover, the structural stability of the STR-211+2-NA complex as revealed by molecular dynamics analyses further confirms the role of STR-211 as a receptor for 2-NA recognition. By employing well-established assays for odorant perception such as chemotaxis, aversion response and lawn avoidance assay, we have provided the *in vivo* experimental evidence for the crucial role of STR-211 receptor in mediating 2-NA perception. Collectively, these findings imply that STR-211 is the candidate olfactory receptor for 2-NA recognition in *C. elegans* and highlight the importance of integration of computational methods with *in vivo* behavioral analysis in deciphering the ligand-receptor interactions and experimental validation. This study, along with an unpublished study (Jin et al., 2026), are the first to identify two distinct GPCRs required for 2-NA sensing.

In the next set of experiments, we have demonstrated that the 2-NA induced freezing response is regulated by STR-211 receptor. We discovered that the receptor STR-211 is contributing to the immediate behavioral phenotype of freezing response in worms such as immobility and halted feeding produced by 2-NA exposure. However, the receptor does not affect the 2-NA stress induced long-term effects such as decrease in the number of progenies. This implies that immediate and long-term health consequences of fear induced stress might be regulated by two distinct signalling mechanisms. Here we are not ignoring the possibility of the role of other chemosensory neurons and receptors expressed in these neurons in regulating the 2-NA induced fear-like behavior. We can expect a combination of receptors and neurons involved in mediating the 2-NA odor evoked freezing following the basic principle of combinatorial coding in the olfactory system. STR-211 is expressed in ASJ and AIY amphid sensory neurons. It will be interesting to find out if STR-211 is working through one or both of these neurons for regulating the freezing response in worms. Understanding the molecular basis of pathogen associated compounds detection and associated behavior in *C. elegans* has broader implications. Many aspects of chemosensory signaling and innate behavioral responses are evolutionarily conserved, offering insights into how more complex organisms interpret environmental threats. Our study demonstrates a robust, simple, cost-effective, and environmentally relevant animal model to mimic fear-like behavior often associated with pathological anxiety. Anxiety disorders are caused by a complex mix of genetic, biological, and environmental factors. Genetic amenability of *C. elegans* combined with advantages of the proposed model may be useful for understanding of the neurobiological mechanisms causing anxiety.

## Acknowledgements

We would like to acknowledge Dr. Abhishek Bhattacharya, NCBS, Bengaluru, for providing RNAi clones, and Dr. Jogender Singh, IISER Mohali, for providing the MAH677 *C. elegans* strain.

## Supplementary material

The following is the Supplementary data to this article: **Supplementary Information**

## Author Contributions

All authors contributed to the study conception and design. Material preparation, data collection and analysis for all *C. elegans* experiments were performed by Anubhuti Dixit, Anisha Bhola and Avachi Azad. Material preparation, data collection and analysis for in silico experiments were performed by Tithi Thakur and Hina Bansal. The first draft of the manuscript was written by Anubhuti Dixit and all authors commented on previous versions of the manuscript. All authors read and approved the final manuscript.”

## Data availability

The datasets generated during and/or analysed during the current study are available from the corresponding author on reasonable request

