## Supplemental Information for "A pathogen-associated odorant induces fear-like response regulated by an olfactory receptor STR-211 in *Caenorhabditis elegans*"

**Figure S1:**

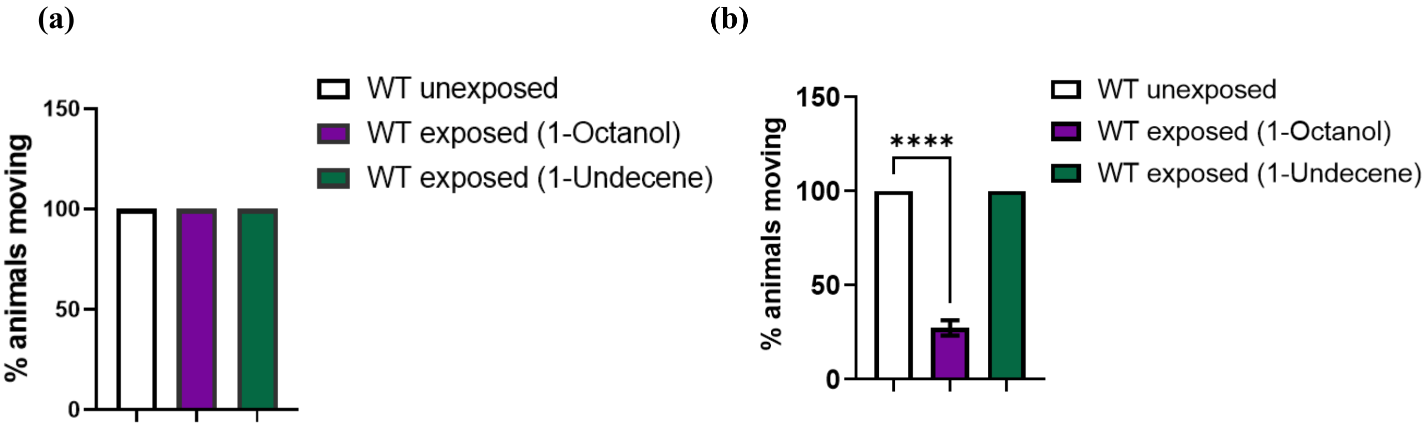


**Figure S1:** Freezing response assessment in C. elegans. (a) Worms were exposed to 3ul of pathogen associated volatile compounds 1- octanol and 1-undecene and percent animals moving were calculated after 4 minutes of odor exposure. (b) Worms were exposed to 6 ul of pathogen associated volatile compounds 1- octanol and 1-undecene and percent animals moving were calculated after 8 minutes of odor exposure. ****=p<0.0001

**Figure S2:**

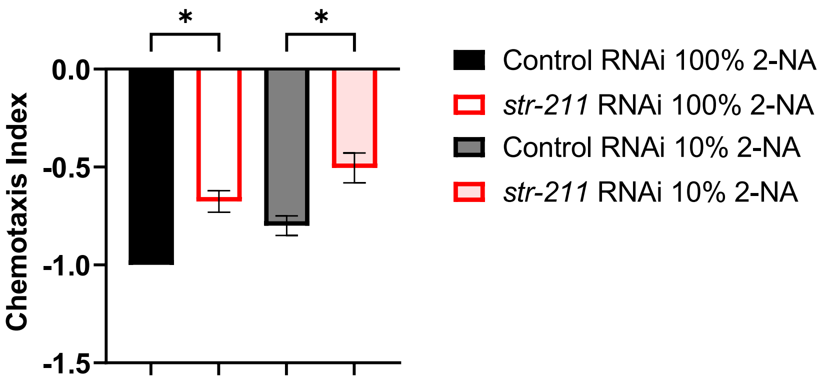


**Figure S2:** Chemotaxis assay of control and str-211 RNAi worms against 100% (undiluted) and 10% (1:10 diluted) concentrations of 2-NA. *=p<0.05

**Figure S3**

**(a)** **(b)**

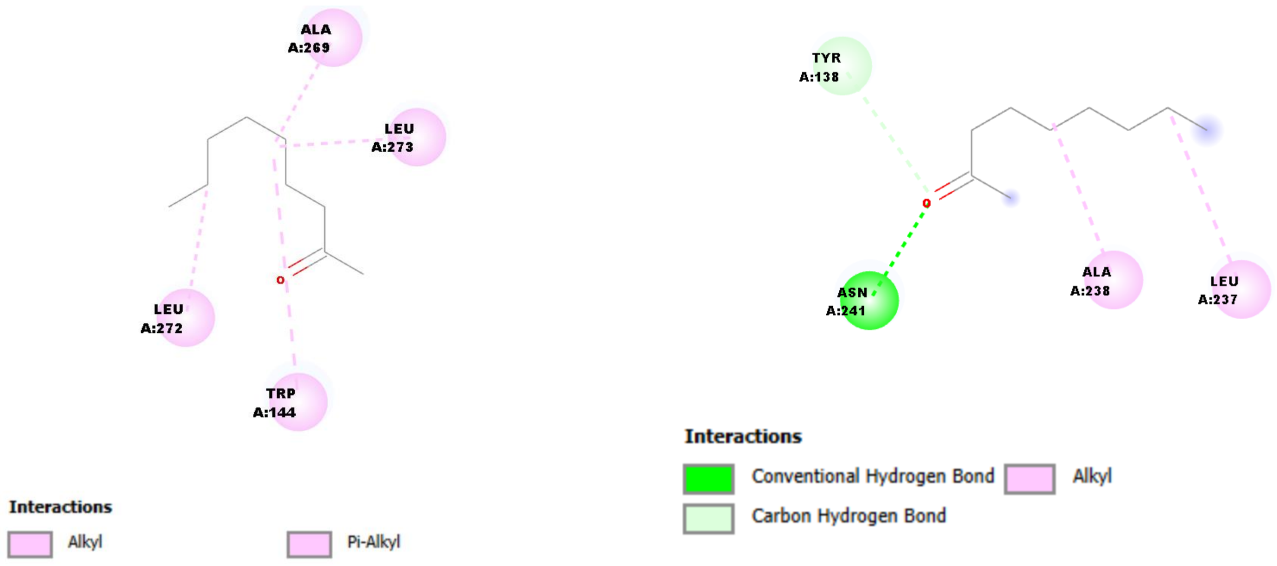


**Figure S3:** Two-dimensional protein–ligand interaction maps of 2-nonanone with the chemosensory receptors (A) STR-92 and (B) SRA-20, illustrating the residue-level interactions predicted from molecular docking.

**Figure S4**

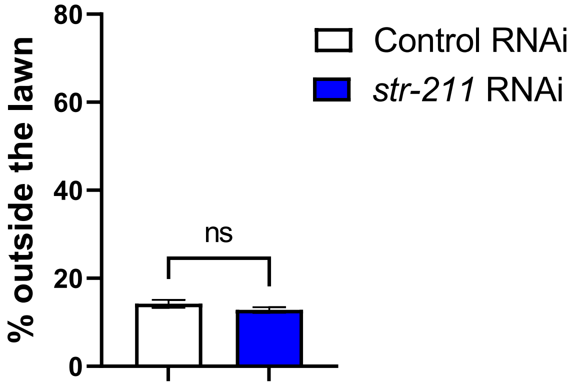


**Figure S4:** Lawn occupancy of control and str-211 RNAi worms in the absence of 2-NA. More than 80% of worms in both the groups were found to be present inside the lawn. ns=non-significant.

**Table 1. Structural validation statistics of the AlphaFold-predicted chemosensory receptors STR-211, STR-92, and SRA-20**

| Parameter | STR-211 (O16439) | STR-92 (O62279) | SRA-20 (O17844) |
|---|---|---|---|
| **Protein length (aa)** | 349 | 343 | 339 |
| **ERRAT Overall Quality Factor** | **99.70** | **93.88** | **93.66** |
| **Ramachandran plot – Most favoured regions (%)** | 95.6 | 94.9 | 96.2 |
| **Ramachandran plot – Additional allowed regions (%)** | 3.4 | 4.7 | 3.8 |
| **Ramachandran plot – Generously allowed regions (%)** | 0.6 | 0.3 | 0.0 |
| **Ramachandran plot – Disallowed regions (%)** | 0.3 | 0.0 | 0.0 |
| **VERIFY3D assessment** | Below threshold[#] | Below threshold[#] | Below threshold[#] |
| **Overall structural quality** | Excellent | Good | Good |

### VERIFY3D scores were below the conventional threshold for globular proteins; however, such behaviour is commonly observed for AlphaFold-predicted transmembrane receptor structures and therefore does not preclude their use in downstream structural analyses.

**Table 2 Binding-pocket characteristics of the selected receptors**

| Receptor | UniProt ID | Pocket Rank | Pocket Score | Probability | No. of Residues | Pocket Center (x, y, z) |
|---|---|---|---|---|---|---|
| STR-211 | O16439 | 1 | 31.64 | 0.899 | 23 | (0.450, −2.312, −11.115) |
| STR-92 | O62279 | 1 | 39.17 | 0.936 | 22 | (5.4179, 4.6722, −3.2418) |
| SRA-20 | O17844 | 2 | 3.59 | 0.117 | 11 | (−13.6055, −7.2022, 20.9794) |
